# An optogenetic assay of *Drosophila* larval motor neuron performance *in vivo*

**DOI:** 10.1101/2025.08.18.670936

**Authors:** Yosuf Arab, Gabriel G. Bonassi, Gregory T. Macleod

## Abstract

**Background:** Over fifty million people worldwide currently live with neurodegenerative diseases, many of which are the result of pathogenic gene variants. Genetically malleable model organisms provide an avenue for research into the genetic bases of these diseases, and the large motor neurons of fruit fly larvae provide a test bed for investigating neuronal mechanisms impacted by pathogenic gene variants. However, it is challenging to collect information from these neurons under physiological conditions as they terminate on muscle fibers that are perpetually contracting - driven by motor neuron burst-firing.

**New Method:** As a test of *in vivo* neuronal performance, we expressed light-activated opsins in motor-neurons of unrestrained intact *Drosophila* larva and used light pulses to drive cyclical body-wall contractions that were captured on camera and analyzed offline.

**Results:** We describe the assembly of an apparatus to systematically activate motor-neurons in *Drosophila* larvae and an image acquisition system to capture the resulting body-wall contractions. To test the utility of the assay we performed a motor-neuron specific knock-down of dMiro, an adaptor for mitochondrial transport into motor-neuron terminals. As predicted, contractions were poorly sustained in larvae with impaired axonal transport of mitochondria.

**Comparison with Existing Methods:** This *in vivo* assay allows for a test of sustained neuronal performance while sidestepping the shortcomings of electrophysiological assays of neurotransmission *in situ* where hemolymph-like solutions may not recapitulate hemolymph properties, axons are severed and where recordings are mechanically disrupted at endogenous firing rates. Secondly, unlike adult climbing assays and larval locomotion assays, performance is assayed independently of the organism’s motivation to perform or ability to detect stimuli.

**Conclusions:** We demonstrated that *Drosophila* 3^rd^ instar larvae cannot sustain body-wall contractions if mitochondria are not delivered to motor nerve terminals - validating a motor neuron performance assay in a model organism suited for molecular genetic analysis.

## 1. Introduction

Neurodegenerative diseases currently diminish the quality of life of over fifty million people worldwide, with disease prevalence increasing as the population ages (GBD 2021 Nervous System Disorders Collaborators, 2024; Imam et al., 2025). A substantial proportion of neurodegenerative diseases can be attributed to pathogenic variants of certain genes, although the proportion varies according to the disease: 100% in Huntington’s disease, 80% in familial Amyotrophic Lateral Sclerosis (ALS) and 5-10% in non-familial (sporadic) forms of ALS, and 30% in Parkinson’s disease (The Huntington’s Disease Collaborative Research Group, 1993; Klein & Westenberger, 2012; Van Damme et al., 2017). The list of known pathogenic variants continues to grow, and the large numbers of familial cases not associated with a known pathogenic variant suggest that many more disease-causing mutations remain to be discovered (Cacace et al., 2016). Genetically-malleable model organisms have a lot to offer in the research of neurodegenerative diseases as they can help elucidate disease processes associated with pathogenic gene variants and provide a platform for the development of therapeutic approaches that aim to improve quality of life (Fig. 1).

**Figure 1.**
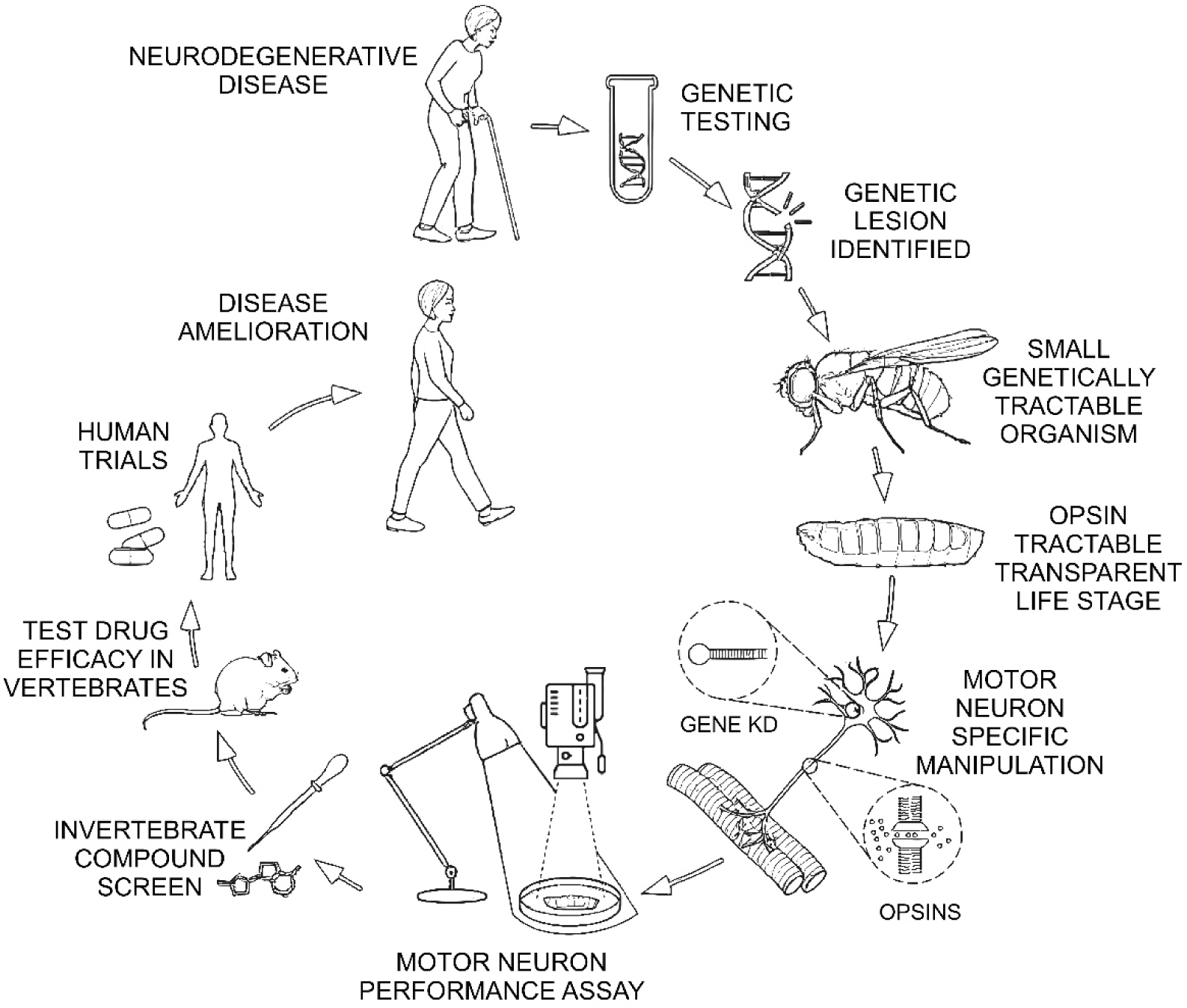
*Drosophila* is a highly tractable model for elucidating the genetic basis of neurodegenerative disease and developing therapies. In this informational age the list of gene variants associated with neurodegenerative disorders continues to grow. Invertebrates such as *Drosophila* provide malleable genetic models in which to elucidate the role of these variants in the pathogenesis and progression of disease, and to develop high-throughput assays that inform therapeutic approaches that might ameliorate the debilitating effects of disease. Figure partially assembled in BioRender then rendered in Artlist.io.

The fruit fly, *Drosophila melanogaster*, is an invaluable model, having been used in numerous studies of neurodegenerative disease (Feany & Bender, 2000; Li et al., 2010; Ma et al., 2023) and helped identify new agents with therapeutic potential (Sanz et al., 2021). It shares approximately 75% of human disease related genes (Reiter et al., 2001), combines high genetic tractability with a well-curated collection of genetic and anatomical data (Milyaev et al., 2012; Ozturk-Colak et al., 2024), and is supported by large repositories of molecular and genetic resources (Thurmond et al., 2019; Ugur et al., 2016). Motor neurons in the larval stage provide ready access for investigating cellular and synaptic mechanisms relevant to many different neuron types not just motor neurons, such as the transport and distribution of mitochondria and dense-cored vesicles (Russo et al., 2009; Tao et al., 2017), axonal regeneration (Hao & Collins, 2017), presynaptic bioenergetics (Chouhan et al., 2012; Justs et al., 2022), and trans-synaptic signaling (Frank et al., 2020), to mention only a few. However, a drawback of working on motor neurons *per se*, is the difficulty of assessing the limits of their performance under endogenous conditions, a highly relevant parameter in neurodegenerative disease. While electrophysiological techniques are able to measure neurotransmission at the neuromuscular junction (NMJ) under highly controlled conditions (Bykhovskaia & Vasin, 2017; Jan & Jan, 1976), the contractile nature of the neuromuscular junction occludes assessment of motor neuron performance during sustained, high-intensity activity typical of locomotion.

To address this limitation, we developed an *in vivo* assay that enables direct activation of motor neurons that drive body-wall contractions, and allows for assessment of neuromuscular performance. Traditional assays, such as geotaxis (climbing assays) can test endogenous motor neuron function (Gargano et al., 2005; Madabattula et al., 2015; Taylor & Tuxworth, 2019), but these assays are susceptible to the banes of behavioral assays - differences in motivation to perform and differences in the ability to perceive prompts to perform (Devineni & Scaplen, 2021; Sun et al., 2018). In contrast, the current approach circumvents the aforementioned confounds by using Channelrhodopsin (ChR2 H134R) to optogenetically stimulate motor neurons with blue light, as demonstrated previously (Dietzl et al., 2007; Pulver et al., 2009), while allowing the investigator to evoke patterned body-wall contractions. This approach was used to test the reliance of motor neurons on dMiro, the *Drosophila* homolog of human Miro1 (RHOT1), a Rho GTPase, which is required for mitochondrial trafficking along neuronal processes (Babic et al., 2015; Nemani et al., 2018; Russo et al., 2009) and whose pathogenic variants have been implicated in Parkinson’s disease (Berenguer-Escuder et al., 2019; Grossmann et al., 2019). Knock down of dMiro expression in motor neurons using dsRNA demonstrated a deficit in the ability of motor neurons to sustain body-wall contractions. These findings highlight the importance of mitochondrial transport for sustaining neurotransmission and validate the utility of this assay for assessing motor neuron performance *in vivo* in *Drosophila* models of neurodegenerative disease.

## 2. Materials and Methods

### 2.1 Genetics

*Drosophila* fly stocks were raised at 24°C on standard medium [recipe from Bloomington Drosophila Stock Center (BDSC), Bloomington, IN]. BDSC provided two UAS-*dmiro*-RNAi fly lines (#27695 and #43973), the UAS-luciferase control line (#35788) and pan-neuronal driver nSyb-GAL4 (#51635). The Vienna Drosophila Resource Center (VDRC) provided a single UAS-*dmiro*-RNAi fly line (#330334). UAS-TagBFP was a gift from Dr Kenneth Irvine. UAS-mitochondrial-mKate, made in this laboratory, has been described previously (Justs et al., 2023). A UAS-CHR2-H134R line, a gift from Dr. Stefan Pulver, allowed the expression of Channelrhodopsin-2 tagged with mCherry (Pulver et al., 2009). OK371-GAL4 (Mahr & Aberle, 2006), which drives primarily in motor neurons with type-Ib terminals [∼22 in each hemisegment; (Hoang & Chiba, 2001)], was recombined on the same chromosome as UAS-CH2-H134R and used to drive expression of both UAS-CH2-H134R and UAS-*dmiro*-RNAi, where the two copies of the driver and opsin were present along with one copy of the RNAi construct.

### 2.2 Solutions, Dissections and Confocal Microscopy

Filet dissections of female 3^rd^ instar larvae were performed on a Sylgard tablet in chilled Schneider’s insect medium as demonstrated by (Rossano & Macleod, 2007), before transferring to hemolymph-like solution #6 (HL6) containing CaCl_2_ added to 0.2 mM (Macleod et al., 2002). Dissected larvae were imaged with a Nikon 60X, 1.20 NA, Plan Apochromat VC water-immersion objective on a Nikon A1R confocal laser scanning microscope fitted with GaAsP detectors. Preparations were scanned sequentially, with the longer wavelength (561 nm) for mKate scanned before the shorter wavelength for TagBFP (405 nm). Images were taken using the same settings and each image presented here (Fig. 6) represents a collapsed Z-series encompassing the full depth of MN13-Ib terminal boutons (between 3 and 5 sections at a 1 μm step size).

### 2.3 Fluorescence Microscopy Data Analysis

Fluorescence images were analyzed using ImageJ software. Z-series of images containing the extent of MN13-Ib terminals were collapsed (average intensity) to generate a single image. Analyses were performed on 16bit images. ROIs were placed around several MN13-Ib boutons to measure the average fluorescence of either TagBFP in the cytosol, or, mKate in mitochondria. Care was taken to avoid any aggregates seen in either TagBFP or mKate. Average fluorescence measurements taken from ROIs placed on the background adjacent to the terminals were subtracted from the bouton ROI measurements to generate a final fluorescence measurement.

Mitochondrial content was calculated as the mitochondrial-mKate fluorescence divided by the cytosolic-TagBFP fluorescence.

### 2.4 Components of the Apparatus

The apparatus consisted of six primary components (Fig. 2A): a red Light Emitting Diode (LED) circuit (Fig. 2A-C), a blue LED circuit (Fig. 2A,D-E), a camera, and an agar-filled Petri dish (agar plate), all on top of a thick aluminum plate. A PC controlled LED circuits to either continually illuminate the larvae on the agar-plate for the benefit of the camera (red LEDs) or excite the motor neurons to fire (blue LEDs).

**Figure 2.**
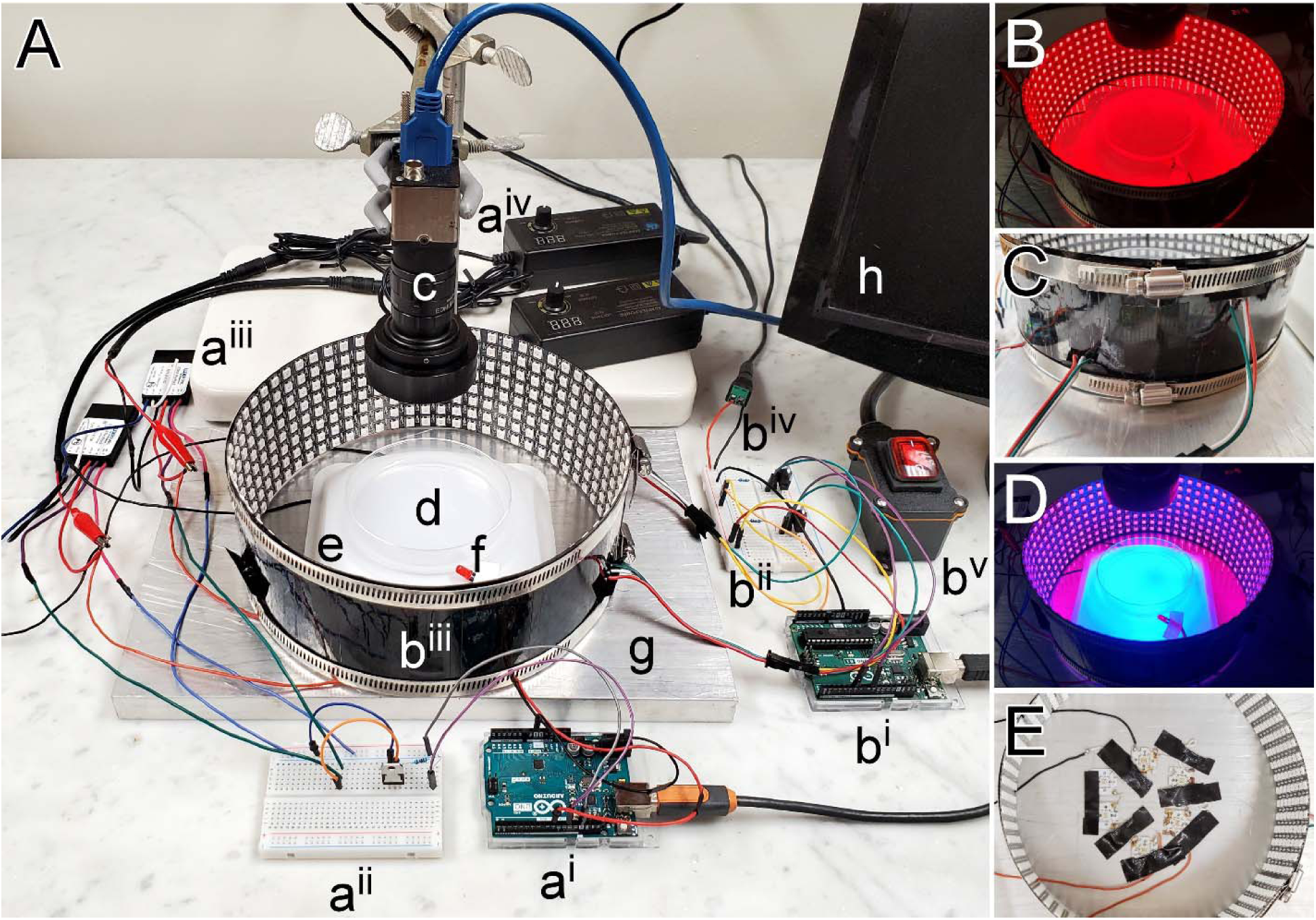
Motor neuron performance assay apparatus. A, Apparatus photograph with wires teased apart to enable circuit connectivity verification; a - blue LED circuit with an Arduino board (a^i^) and separate breadboard (a^ii^), two LED drivers (a^iii^), two 10V power supplies (a^iv^); b - red LED circuit with an Arduino board (b^i^) and separate breadboard (b^ii^), two LED matrices joined in a cylinder (b^iii^), beadboard connection to a 5V power supply (out of view) (b^iv^), inline cord switch for on/off power control of LED matrices (b^v^); c – camera with a blue light filter in front of the lens (out of view); d - agar-filled Petri dish; e - inverted weigh-boat obscuring the blue LEDs beneath; f - single red LED; g – 19 mm thick aluminum plate heat-sink; h - computer used to control the Arduinos through our open-source Bonsai program. B, LED matrices programmed to emit red light. C, Detail of the side of the RGB matrices joined in a cylinder by worm-gear duct-clamps. D, Blue light emitting from LEDs beneath an inverted weigh-boat used to excite opsins within the larvae, programmed to switch on intermittently, while the surrounding LED matrices emit red line continuously so that larva can be observed without interruption. E, Blue LEDs (two strings of 3 each) fixed to the aluminum plate in a hexagonal pattern with thermal paste and tape. Weigh-boat and insulation removed to reveal detail.

### 2.5 Red LED Assembly

Ambient lighting was provided by two addressable RGB LED panels (80 x 320 mm; WS2812B matrix) which surrounded the agar-plate. The LED panels were taped together in a short cylinder which was placed on its end to surround the agar-plate (Fig. 2A-C). The panels were connected in parallel to a 5V power supply (Alitove, ALT-0503) and an Arduino was used to program the LED circuits (Fig. 3A).

**Figure 3.**
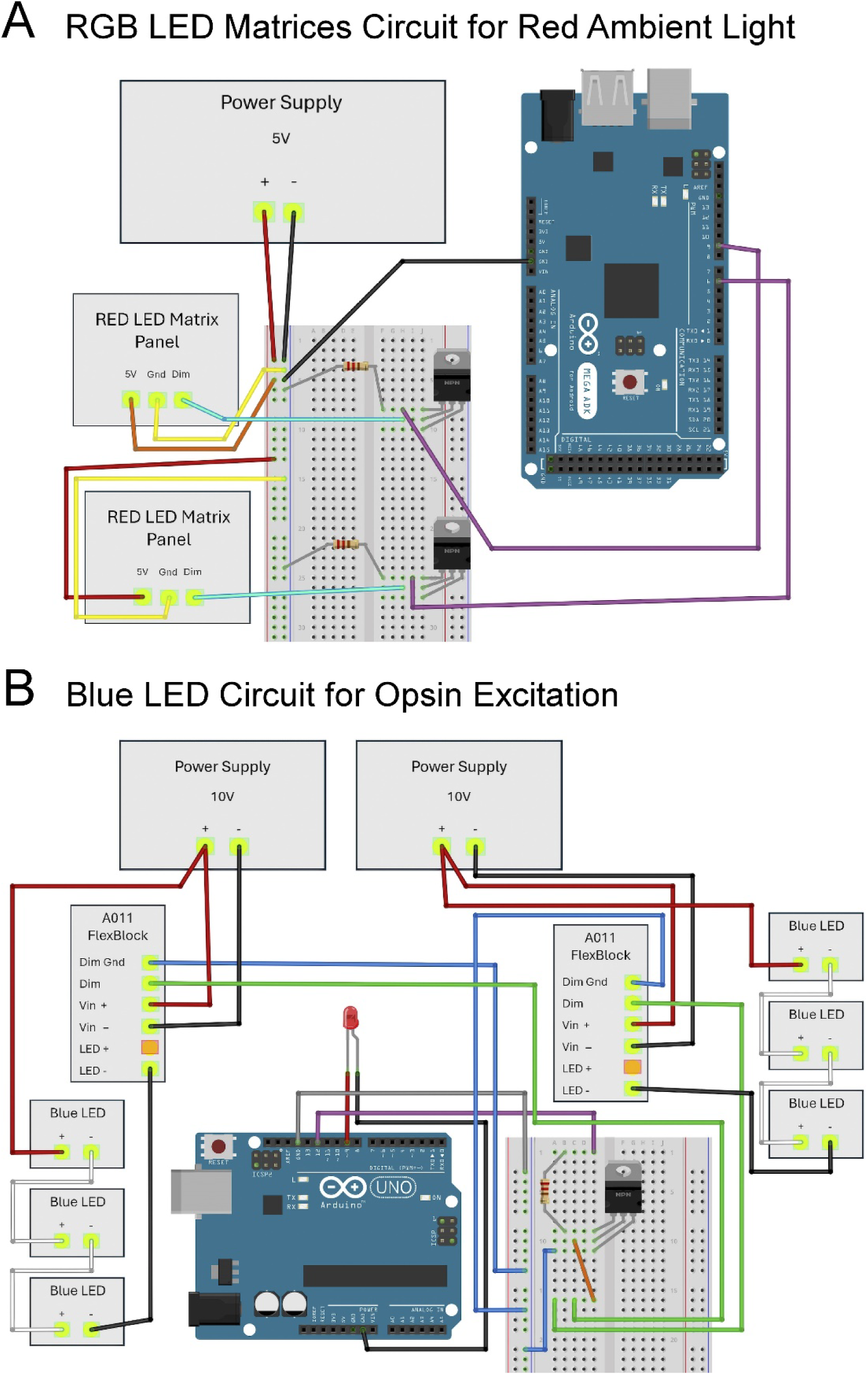
Schematic of the circuits controlling blue (excitation) LED arrays and red (ambient) LED arrays. A. Schematic of the wiring configuration for two RGB LED matrix panels. Matrix panels were wired in parallel between the positive pole of a 5V power supply and the base (pin 1) of a dedicated NPN Darlington transistor (TIP125). The data input pole (DIN) of each matrix panel was connected to the collector (pin 2) of its respective transistor. Connections were made from the base (pin 1) of each transistor to digital pins 9 and 6, respectively, on the Arduino, via 10 KOhm resistors. B. Schematic of the wiring configuration for 6 blue LEDs and a single red LED. Two groups of blue LEDs are depicted, wired in series, with each group having a dedicated A011 FlexBlock driver and an adjustable 10V power supply. One end of each group of blue LEDs connects to the positive pole of one of two power supplies and the other end connects to the LED negative pole of one of two drivers. Connections were made between the dimmer ground (Dim Gnd) on each FlexBlock to a common ground on the breadboard, while another connection was made from each FlexBlock dimmer pole (Dim) to the collector (pin 2) of an NPN Darlington transistor (TIP120). A 10 KOhm resistor was placed between the base (pin 1) of the transistor and digital pin 9 of an Arduino Uno. The emitter (pin 3) of the transistor was connected to the breadboard common ground. Additionally, a single red LED diode was wired between digital pin 12 and ground of the Arduino Uno. Schematics in A and B assembled in Fritzing.

### 2.6 Blue Light Assembly

To excite the motor neurons we used 6 blue LEDs (Luxeon Star, SZ-05-H3; 470nm). Two groups of 3 LEDs were connected in series and arranged hexagonally with approximately 15 mm between each (Fig. 2E). LEDs were fixed to an aluminum plate (200 x 300 x 19 mm) with thermal paste to dissipate heat and taped in place. An inverted weigh-boat (140 × 140 × 22 mm; Heathrow Scientific, HS1421C) was placed over the LEDs, and the agar-plate was placed directly on the weight boat. To provide sufficient power, each group of 3 LEDs was connected to a dedicated 700mA LED driver (Luxeon Star, A011-D-V-700) and a 10V adjustable power supply (Kejiesheng, KJS-1509) (Fig. 3B). Each LED driver was connected to a half-size breadboard containing a Bojack Darlington transistor (TIP120) and a 10 KOhm resistor. An Arduino Uno interfaced with the breadboard and was connected to a PC computer using an Arduino USB 2.0 Cable Type A/B, thus providing the capability to program the blue LEDs to alternate between on and off. Lastly, a single red LED diode was wired to the same Arduino Uno to mirror the status of the blue LEDs. The thermocouple probe of a K-type thermometer (Fisher Scientific; 15-078-38) was used to measure the temperature on the top surface of the agar, over the course of a 20-minute experiment, which did not rise by more than 1°C.

### 2.7 Camera

The camera (Edmund Optics, 1312M 1/2“ CCD Monochrome USB) was situated approximately 190 mm above the agar-plate using a ring stand. It was fitted with a fixed focal-length lens (Edmund Optics, 33-301; 6mm/F1.85) and an M43-x0.75 adapted housing to fit a 50 mm diameter blue light long-pass filter (Edmund Optics, 18-884; 640nm cut-on). The filter prevented the camera from detecting changes in the blue LEDs but allowed detection of synchronous changes in the single red LED. A blackout curtain was placed over the entire apparatus to block external light.

### 2.8 Preparation of Agar-Filled Petri Dishes

Preparation of a 2% agar-filled Petri dish (agar plate) began with 0.8 grams of agarose (Apex, 20-101) dissolved in 40 mL of distilled water. It was transferred to a 250 mL glass beaker and microwaved on high for 1.5 minutes, with a swirl every 30 seconds. The solution was reduced to 27 mL and divided evenly among two 88 mm Petri dishes, covered, then refrigerated at 4°C until hardened.

### 2.9 Preparation of All-Trans-Retinal Food

Flies were raised in food vials containing All-Trans-Retinal (ATR; Sigma-Aldrich, R2500-1G), a molecule necessary for the proper folding of Channelrhodopsin-2. 1 mL of distilled water was added to roughly 50 mL of yeast fly food, mixed, microwaved for 25 seconds, then mixed again until it was a homogenous mixture. Once cooled to a lukewarm temperature, 250 µL of 100 mM ATR in ethanol stock was mixed into the liquid fly food (∼500 µM final ATR concentration; 0.5% ethanol). It was then evenly divided among six vials and left to cool and solidify in a dark box. After solidification, the flies were transferred into the ATR food vials and stored in the dark box at room temperature.

### 2.10 Arduino Setup

Arduinos are inexpensive single-board microcontrollers that can be connected to a computer via USB for programming purposes. Two Arduino boards were used to coordinate an alternating blue LED arrays (and an accompanying single red LED) and a set of two constantly illuminated red LED panels. Programming for both light systems was done in the Arduino Integrated Development Environment (IDE; Version 2.2.1; code attached). One Arduino board switched the blue LEDs on and off according to a prescribed protocol while mirroring this protocol in a single red LED. The blue-LED Arduino program “Blue_LED_Light_Program” was uploaded then initiated by starting our open-source Bonsai program; both programs can be found in the supplemental data section. The second Arduino controlled the red LED panels, independently of Bonsai, through Arduino IDE software running “Red_LED_Matrix_Program”, which can be found in the supplemental data section. This required a library, named “Adafruit NeoPixel,” to be installed via the “Manage Libraries” facility under the “Include Libraries” tab from the “Sketch” menu. The software must be restarted after putting the files in place and the program must be uploaded to the Arduino by selecting the arrow on the top left corner of the program file. At the end of any trial, the Bonsai program can be terminated via a stop button, and the blue LEDs can be deactivated via the blue LED Arduino’s reset button. Power to the red LEDs can be terminated independently using an inline cord switch. All supplemental data, including.bonsai and.io files, can be found in this public GitHub repository: https://github.com/GabrielBonassi77/Motor-Performance-Assay

### 2.11 Bonsai Pipeline Setup and Configuration

The data pipeline (Fig. 4) was created using a visual programming language; <u>Bonsai 2.8.1</u> (.zip). The Bonsai Starter Pack was installed from the tools tab utilizing the search bar. Our open-source Bonsai program, titled **“**BioThresh”, can be found in the supplemental data section.

**Figure 4.**
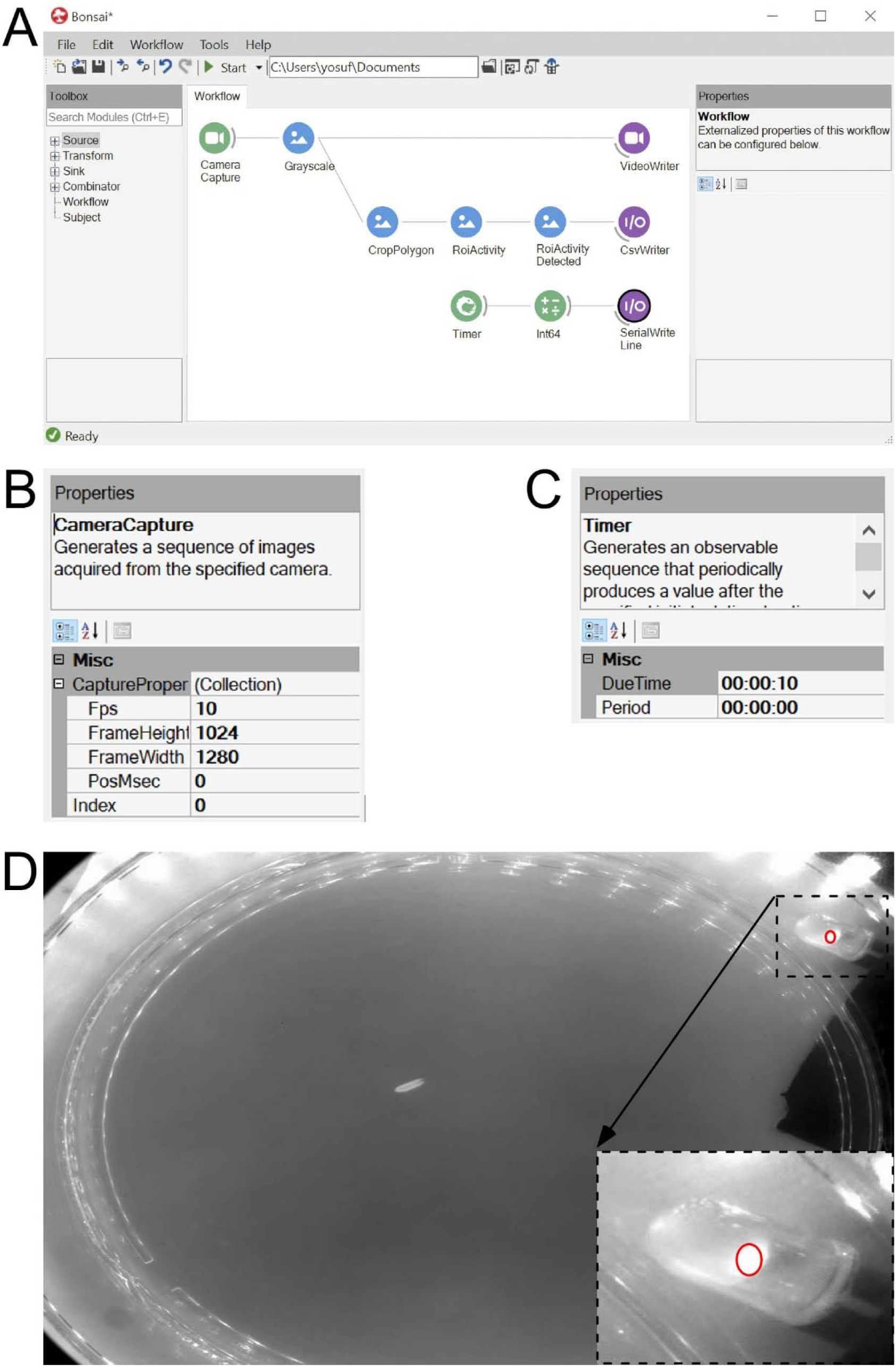
Software Interface for Blue LED Control and Data Collection. A. Depiction of two pipelines within a workflow file [Biological Thresholder (BioThresh)] of the Bonsai program. The top portion of the first pipeline (first row) was used to record a video of a larva using the camera. See single frame captured in D. The bottom portion of the first pipeline (second row) was used to create a Region of Interest (ROI; CropPolygon node) and place it on the red LED (inset in D) that is powered by the same circuit that powers the blue LEDs. This allows for a record of pixel intensity in the ROI and thus a record to corroborate when the blue LEDs were on. The second pipeline (third row) was used to control the Arduino program that supplied power to the blue LEDs (Blue_LED_Light_Program). B. Settings (Properties and Miscellaneous dialogue boxes) used for the “Camera Capture” node in the first pipeline. C. Settings for the “Timer” node in the second pipeline. D. An image of a larva within the center of an agar plate illuminated with red LED matrices visible in reflection. Inset: a small ROI placed over the single red LED that illuminates in synchrony with the blue LEDs.

### 2.12 Camera Setup

The VideoWriter and CsvWriter nodes were disabled prior to testing. In the “Camera Capture” node, the three dots adjacent to “(Collection)” were selected and the settings and values were added as shown in Figure 4. If a video was previously recorded, a pipeline identical to the first pipeline (Fig. 4A & 4B) was created, but the Camera Capture node was replaced with a FileCapture node. A camera was selected from the index tab, and the “Start” button was clicked to run the Bonsai program. To view the camera’s live video stream, the camera “Camera Capture” icon was double clicked.

### 2.13 Collection of Blue LED Status Data

To synchronize the start button of the Bonsai program and the blue LED Arduino program, the corresponding port number to the blue LED Arduino circuit was typed in the “Serial Write Line” node. The timer node’s “due time” setting was used to delay the blue LED program by 10 seconds to obtain a baseline of the larva prior to stimulated contractions (Fig. 4A). Data pertinent to the blue light status, i.e. whether the blue LEDs were on or off in each frame, were collected using the nodes in the bottom portion of the first pipeline. In the “CropPolygon” node, the three dots were selected in the box adjacent to the “Regions” setting. An ROI was created by clicking and dragging (press shift if you want the ROI to be elliptical), and it was placed atop the red LED (Fig. 4D).

Only one ROI was created and any others were deleted. This process was repeated for the “ROIActivity” node and the “Operation” setting was changed to “average”. The “ROIActivity Detected” node was used to see if the blue LED status was properly depicted. The threshold value was adjusted until consistent accuracy, reflective of the blue light status, was achieved. When recorded to a.csv file, information on the blue LED status was provided for each frame.

### 2.14 Running Trials

Laser safety glasses should be worn to protect against these particularly bright LEDs. The red LED matrices were turned on and the blue LED code was uploaded to the respective Arduino circuit. The agar plate was cleaned with a wet Kimwipe, then placed on the inverted weigh-boat under the camera. Under the VideoWriter node, we selected the ellipsis (…) next to the FileName and created a new file with the.avi extension. A corresponding file name was also made under the CsvWriter, then both nodes were then enabled. The two power supplies connected to the blue LEDs were set to 10V each. The larva was placed in the agar plate, and the program was initiated. The camera recorded the larvae at 10 frames per second. The program, blue LED alternation, was programmed to begin running after 10 seconds of recording a baseline. The trials ran for 20 minutes after the baseline with the blue LEDs alternating between 2 seconds on and 1 second off. In-between trials, the agar plate was wiped with a wet Kimwipe to clean and prevent the agar from drying.

### 2.15 Fiji/ImageJ/WrmTrck Data Analysis

ImageJ was used to process the.avi video files and the WrmTrck plugin was downloaded to analyze larva contractions. The video was uploaded as a virtual stack and converted to grayscale (Fig. 5A). The video was cropped to contain the entire track of the larva throughout the video. The ‘Gaussian Blur’ filter (radius=1) was applied to the video (Fig. 5B). The larva was isolated using the ‘Subtract Background’ option with 50 pixels rolling ball radius (Fig. 5C). A binary version was created by utilizing the ‘Threshold’ tab and Otsu option (Fig. 5D). The parameters were adjusted to highlight the larva against the background. Once a binary video of a black larva on a white background was made (Fig. 5E), the WrmTrck plugin was used. The settings were altered to encompass the larva and the raw area/perimeter/distance data were selected (Fig. 5F). Perimeter measurements most reliably reflected larval contraction and were least perturbed by larval “rearing” behavior. The perimeter data derived from the video were then copied and pasted into the corresponding.csv file, then opened in Microsoft Excel and aligned with blue light status data. The data were then plotted against time.

**Figure 5.**
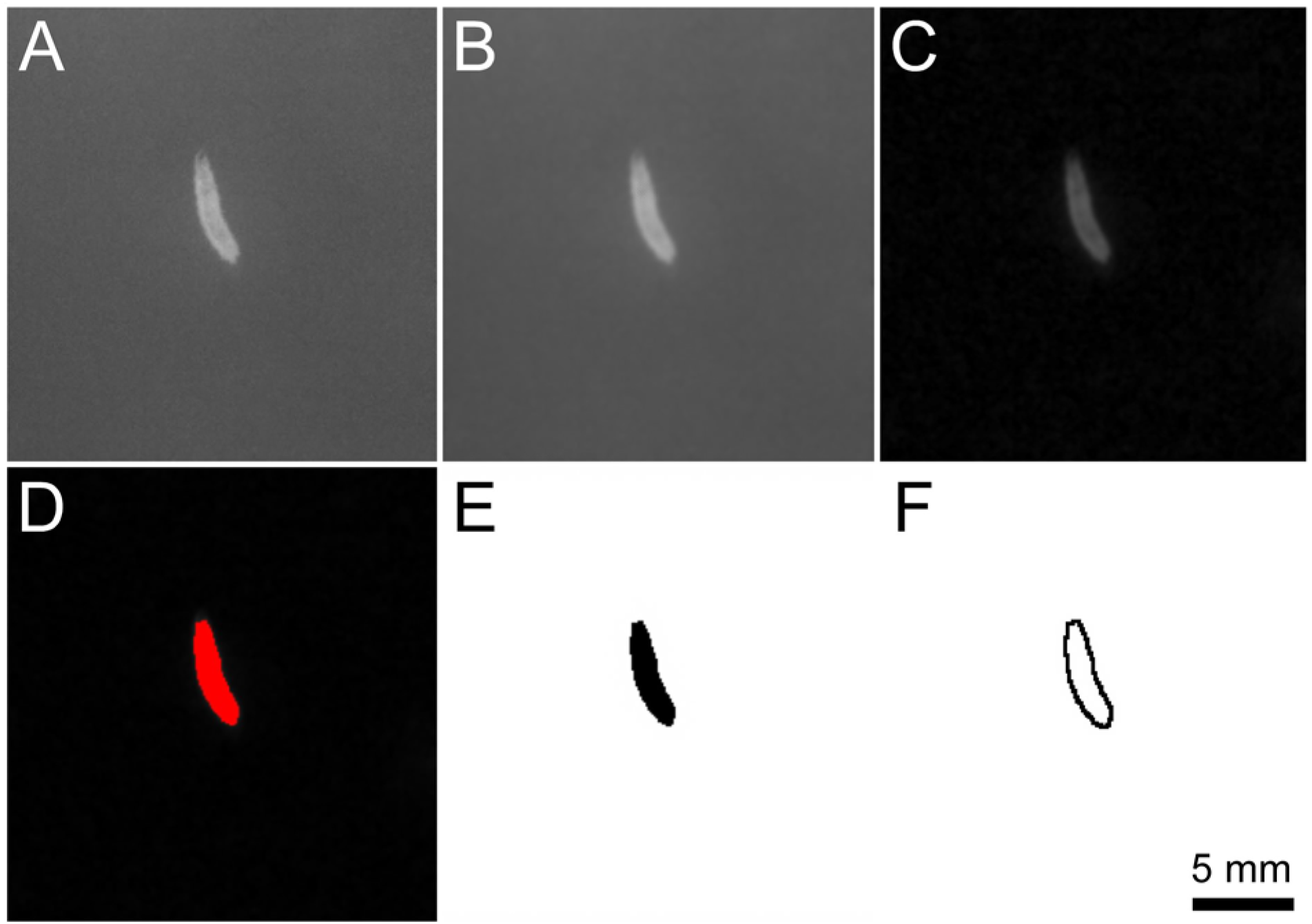
Image processing sequences within ImageJ. A. A raw image of a larva before any video processing. B. Larva after a Gaussian Blur filter was applied. C. Background was removed to isolate the larva from the rest of the video. D. Threshold applied using the Otsu method to highlight the larva. E. Video image converted to a binary video with the larva in black (allows WrmTrck to identify the larva when looking for objects to track). F. Perimeter pixels of the larva. The perimeter was tracked, frame by frame, throughout the video and the perimeter length was plotted against time.

**Figure 6.**
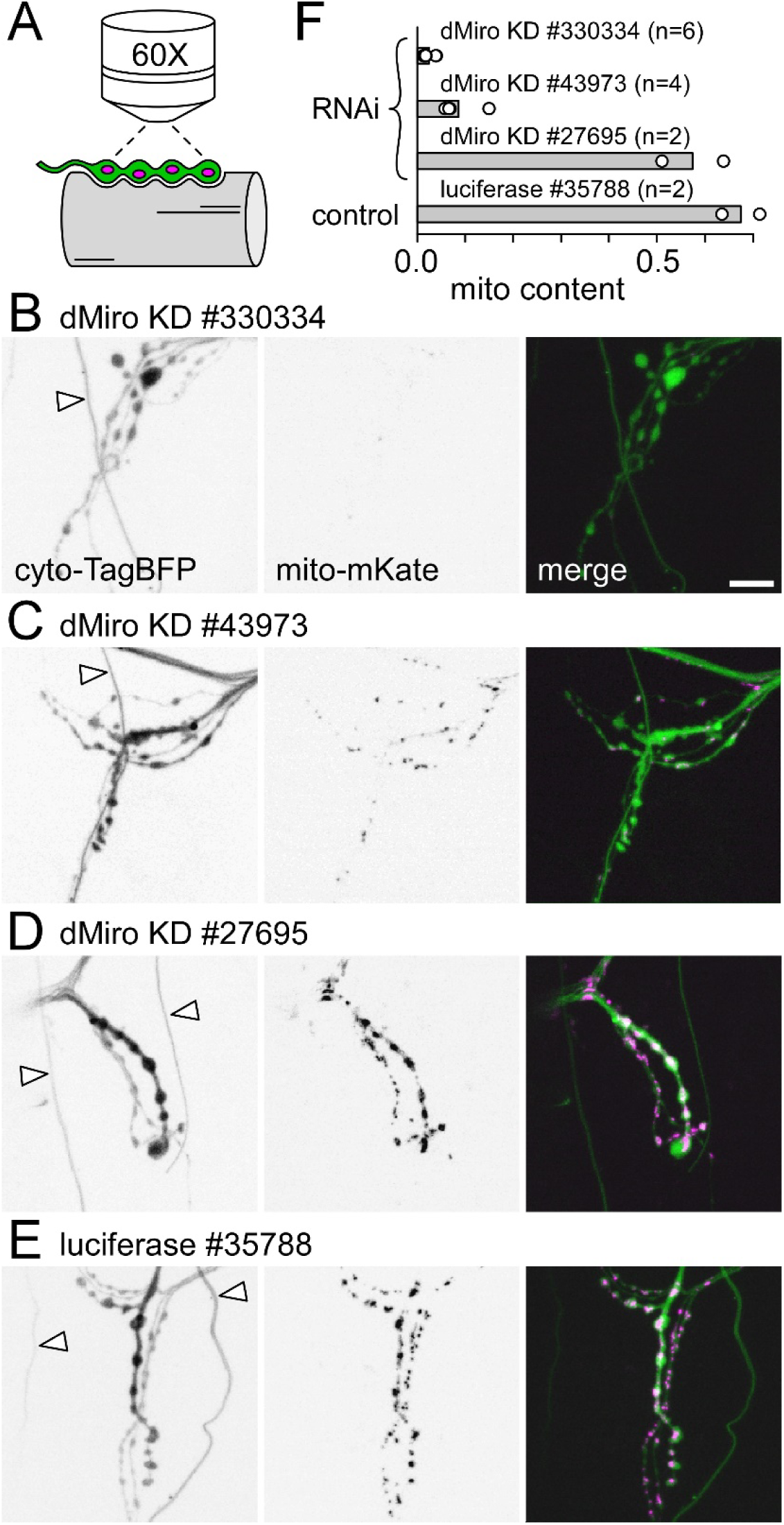
RNAi-mediated knockdown of dMiro diminishes presynaptic mitochondrial content. **A.** A Schematic of a motor neuron terminal on larval body-wall muscle fiber #13 being examined for cytosolic TagBFP fluorescence (green) versus mitochondrial-targeted mKate (magenta) on a confocal microscope. **B-E.** Inverted greyscale images of fluorescence from live terminals on muscle fiber #13. nSyb-GAL4 was used to drive UAS-mitochondrial-mKate and UAS-cytosolic-TagBFP as well as UAS-*dmiro* dsDNA to knock down dMiro (B-D) or UAS-luciferase as a control (E). Stock numbers for each UAS construct fly line are shown. Cytosolic TagBFP is shown in panels to the left, while mitochondrial mKate is shown in the middle column of panels. Images of cytosolic TagBFP, rendered in green, merged with images of mitochondrial mKate rendered in magenta to show relative positioning, in the panels to the right. Fluorescent “traces” denoted with arrowheads in the left panels (B-E) represent autofluorescence from trachea (airways) excited by blue light. Scale bar 10 μm. **F.** A plot of the mitochondrial content in the type-Ib “big” terminals (MN13-Ib) after dMiro was knocked down by the RNAi lines listed. Bar length represents the average of n MN13-Ib terminals in hemi-segment 4. Each n represented by a circle. N (larvae) = n/2.

### 2.16 Data Normalization

The average larval perimeter was calculated immediately before (0-2 seconds) the first blue LED stimulus and used as a baseline. All perimeter values for the trial were then divided by the 0-2 seconds pre-stimulus window values to normalize the perimeter for the duration of the trial, thus reducing inter-larvae differences in size and posture.

## 3. Results

### 3.1 Testing dMiro RNAi reagents

To test the requirement for dMiro in motor neuron performance we knocked down dMiro expression (dMiro KD) in motor neurons alone and monitored the ability of intact larvae to sustain body wall contractions. As a first step, we performed a preliminary screen of the capacity of different RNAi lines to knock down dMiro. dMiro KD will disrupt microtubule-based transport of mitochondria and thereby reduce presynaptic mitochondrial content (Guo et al., 2005) which we assessed here using confocal microscopy to collect images of TagBFP fluorescence in the cytosol and mitochondrial-targeted mKate (Fig. 6A). We used the GAL4/UAS binary expression system to drive dmiro dsRNA expression in larval motor neurons using three different RNAi lines (VDRC #330334; BDSC #43973 and #27695; Fig. 6B-D, respectively). As a control, luciferase expression was driven from a UAS construct in a VALIUM10 vector (BDSC #35788; Fig. 6E) similar to the VALIUM22 vector that harbored most of the UAS-*dmiro* dsRNA constructs. Mitochondrial content was quantified as mitochondrial fluorescence intensity divided by cytosolic fluorescence intensity. The dmiro dsRNA lines showed a wide range of efficacy, with #330334 showing an approximate 96% reduction, #43973 showing an approximate 87% reduction and #27695 showing negligible (∼15%) reduction of mitochondrial content (Fig. 6E). We proceeded to opsin-enabled performance assays with BDSC #43973 as it provided a substantial reduction in mitochondrial content but without the vacuolated muscle fibers and weak larvae associated with the more pronounced dMiro KD of VDRC #330334 expression.

### 3.2 Establishing a test protocol

Our motor neuron performance assay required control of excitation of the very same motor neurons in which dMiro was diminished and this was achieved by driving *H134R-ChR2* and dmiro dsRNA expression in the same neurons (Fig. 7A-B). Our motor neuron stimulation protocol was designed to accomplish several objectives. First, it had to impose a substantial metabolic load, equivalent to locomotion for many minutes.

**Figure 7.**
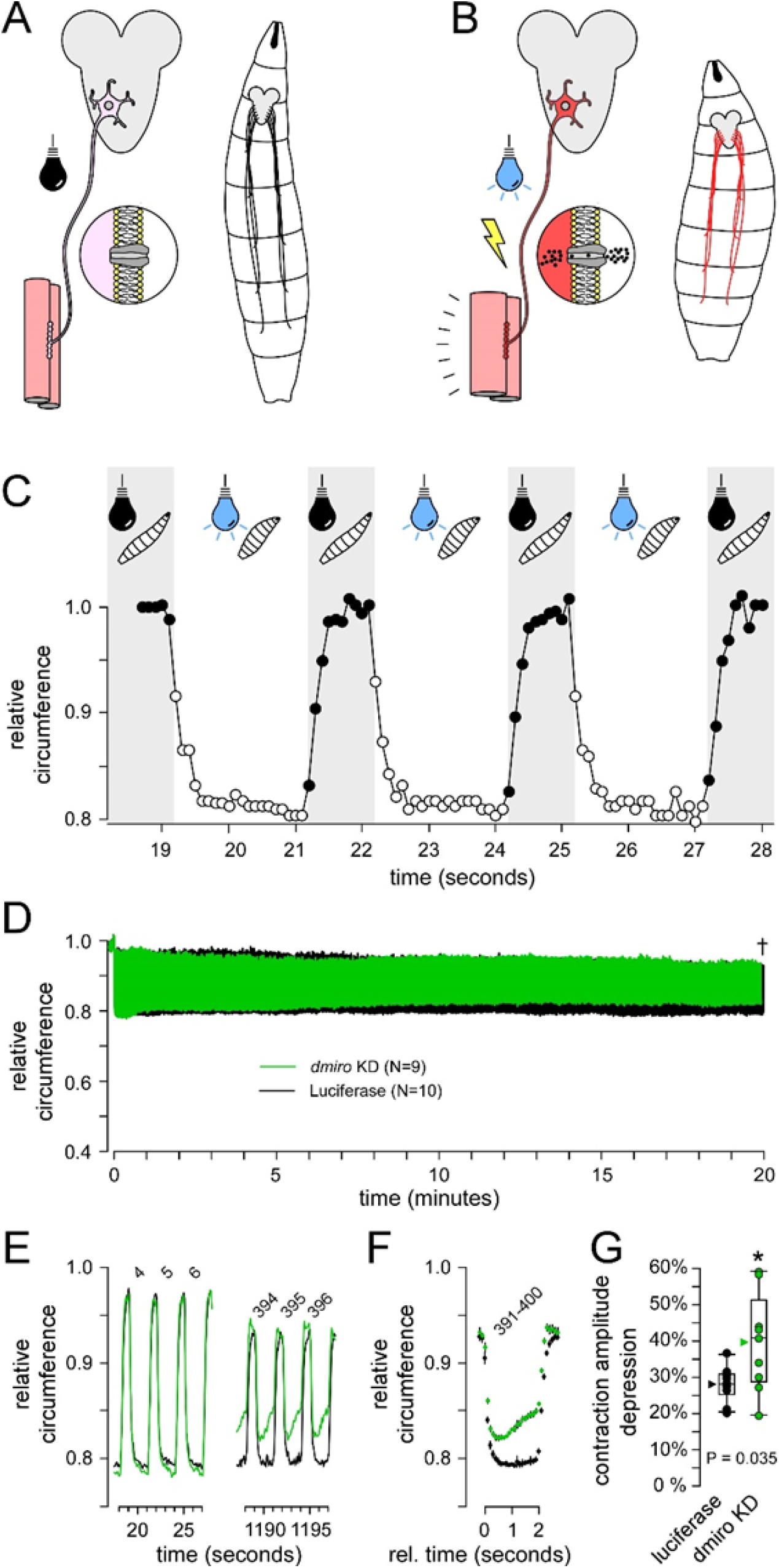
Analysis of musculoskeletal performance after dMiro KD. A. The GAL4/UAS system allows for motor neuron specific expression of Channelrhodopsin-2 (ChR2), a cation-permeant light-activated channel, which is trafficked to the plasma membrane. B. Illumination of *Drosophila* larvae with blue light opens ChR2 depolarizing the plasma membrane causing burst firing, neurotransmitter release and muscle contraction. C. Blue light illumination causes immediate and sustained larval contraction that ceases upon cessation of illumination, and can be repeated for hundreds of cycles. The trace shows the change in the normalized perimeter of a single larva, without data averaging or smoothing, in response to illumination as depicted. D. Plot of cyclical larval contraction showing the average normalized perimeter measurement of individual larvae in which dMiro has been knocked down [dMiro KD; UAS-*dmiro* dsDNA (#43973); N=9] and the control in which the same motor neuron driver (OK371) is used to express luciferase [#35788; N=10]. The dagger indicates that the final 10 seconds of the dMiro KD trace was omitted to allow a comparison between genotypes. E. Plots of 3 cycles (numbered) of larval contraction near the start of the 20 minute regime in D, and near the end. F. A plot of the average of the last 10 contraction cycles (391-400) in D. Standard deviation is shown, almost obscured by the size of the data point. Plots in C-F are normalized to the average perimeter value over a 2 second period immediately before the first illumination cycle. G. Box plots of the amplitude of the change in the normalized perimeter between rest and maximum contraction over the last 3 contraction cycles compared to the first 3. Each point represents one larva; group mean (arrowhead), median (line), 25-75 percentiles box and 10-90% whiskers. A Student’s T-test was used to test for significance.

Second, it had to replicate the cyclical metabolic load that motor neuron’s experience during locomotion. Third, it had to deny unrestrained larvae the opportunity to crawl out of the field-of-view. Lastly, it had to allow signal averaging across multiple larvae and therefore consistency in the protocol from one larva to the next. We settled on a stimulation protocol consisting of 2 seconds of illumination followed by a rest for 1 second (excitation for 67% of the time) (Fig. 7C). During the illumination period, motor neurons will depolarize resulting in burst firing (Pulver et al., 2009). The average duty cycle for motor neurons during fictive locomotion in 3^rd^ instar larvae has previously been established at approximately 78% (Klose et al., 2005). Therefore, if the motor neurons fire at a frequency close to their endogenous firing frequency during opsin-imposed burst firing, the motor neurons would be expected to experience a time-averaged metabolic load of approximately 86% (i.e. 67%/78%) of that experienced during unrestrained locomotion.

### 3.3 dMiro KD larvae show a body-wall contraction deficit

To refine our focus on motor neurons we used the OK371-GAL4 motor neuron driver, rather than the pan-neuronal driver nSyb-GAL4 driver that was used in our preliminary screen. Importantly, two copies of the OK371-GAL4 driver were used to ensure a high level of opsin expression and dMiro KD comparable with a single copy of the stronger pan-neuronal driver. Individual larvae were placed on the agar plate of the apparatus shortly prior to each trial, as described in section 2.14 of the Methods. Larvae were submitted to cyclical illumination for 20 minutes before data were analyzed and pooled (Fig. 7D), as described in sections 2.15 and 2.16 of the Methods. Despite the fat and viscera that surrounds the motor nerves in 3^rd^ instar larvae, blue LED illumination was sufficiently bright to drive body-wall contractions unabated for 20 minutes. dMiro KD larval contraction was indistinguishable from control at commencement (Fig. 7E), but after 20 minutes dMiro KD larvae were unable to sustain contraction over each 2 second period of illumination (Fig. 7E-F), and the magnitude of the dMiro KD contraction was reduced relative to control (P= 0.035; Fig. 7G).

## 4. Discussion

Here, we described an opsin-enabled performance assay for motor neurons *in vivo*. Our knockdown of dMiro resulted in a reduction in presynaptic mitochondrial content by approximately 87%, which might be expected to result in substantial deterioration in motor neuron performance. Our data demonstrate deficits in body-wall contractions, signaling deficits in motor neuron performance, but the contractions were more robust than might have been predicted from the extent of the reduction in mitochondrial content.

Previous studies that demonstrate a reduction in presynaptic mitochondrial content in *Drosophila* larval motor neurons provide context for these findings. Null mutations in *dmiro* can result in an almost complete absence of presynaptic mitochondria (Guo et al., 2005; Russo et al., 2009), and neurotransmission at the NMJs of these mutants fatigues within tens of seconds even when the motor neuron is driven at only 10 Hz. While motor neuron firing rates associated with opsin activation *in vivo* are not known, the extent of body-wall contraction in this study suggests that the firing rates approach the rates that drive locomotion - rates that are thought to exceed 20 Hz, depending on the motor neuron identity (Chouhan et al., 2012). Therefore, it is impressive that these larvae can sustain such a robust level of body wall contraction, and presumably, cycling motor neuron firing well in excess of 10 Hz, for a period of 20 minutes. Hypomorphic mutations in Dynamin related protein 1 (*Drp1*), responsible for mitochondrial fission, result in a reduction of presynaptic mitochondria by approximately 80% (Verstreken et al., 2005) similar to the reductions observed in this study. Yet, neurotransmission in *Drp1^2^* mutants with only an 80% reduction in mitochondria also fatigues within tens of seconds when motor neurons are driven at 10 Hz, albeit at a slower rate than *dmiro* nulls.

It would appear that aspects of the *in vivo* conditions here mitigate against fatigue; aspects such as the endogenous hemolymph bathing the NMJ, rather than an artificial saline, and optimal super-fusion of the NMJ during unrestrained contractions. Additionally, the *in vivo* assay used here does not require motor axons to be severed for the purpose of electrical stimulation and control of axon firing. However, we may also be observing a manifestation of the neuromuscular safety factor where electrophysiological measures of neurotransmission might deteriorate substantially before reaching a threshold that impairs body wall contraction (Marrus & DiAntonio, 2005; Wood & Slater, 2001). Some reconciliation can be found in the locomotor capabilities of the *dmiro* and *Drp1* mutants, as they appear to fit with our expectations based on mitochondrial content, if not the electrophysiological assays. Homozygous *Drp1^2^* mutants with a presynaptic mitochondrial content of ∼20% can locomote and feed through all larval stages and pupate on substrate above the food. Homozygous *dmiro*^B682^ null mutants, on the other hand, with a mitochondrial content of <1%, locomote until the 3^rd^ instar larval stage (the stage being tested in this study) when they become “slim” in body form and immobile on the surface of the food substrate. Thus, although a safety factor should diminish the ability of our assay to discern deficits in motor neuron performance, this assay was nevertheless able to detect a deficit corresponding to a presynaptic mitochondrial content that, at least in *Drp1^2^* mutants, does not otherwise reveal itself in larval behavior.

A particularly valuable aspect of this assay is that it circumvents the confounds of motivation and impaired sensory inputs of behavior-based *in vivo* assays of motor neuron function, such as (in flies) wall climbing. Another valuable aspect is its high-throughput potential - valuable for either elucidating molecular genetic mechanisms or pre-clinical drug screening. Yet, the assay as executed here, does have limitations as some larva climbed up the sides of the agar-plate. However, parameters such as light intensity, pulse duration, and rest duration, can be altered to minimize the opportunity for escape. The simplicity of running trials allows for relatively quick, large-scale data collection, and with the aid of programs able to identify and track multiple larvae in a small arena (Thane et al., 2023), statistically robust data could be collected in much fewer trials.

## Supporting information

https://github.com/GabrielBonassi77/Motor-Performance-Assay

